# Mastitis risk effect on the economic consequences of paratuberculosis control in dairy cattle: A stochastic modeling study

**DOI:** 10.1101/646893

**Authors:** L. J. Verteramo Chiu, L. W. Tauer, Y.T. Gröhn, R. L. Smith

**Affiliations:** Department of Population Medicine and Diagnostic Sciences, Cornell University College of Veterinary Medicine, Ithaca, NY 14850, USA; Charles H. Dyson School of Applied Economics and Management, Cornell SC Johnson Business College, Cornell University, Ithaca, NY 14850, USA; Department of Pathobiology, University of Illinois, College of Veterinary Medicine, Urbana, IL 61802, USA

**Keywords:** paratuberculosis, economic model, disease control, mastitis

## Abstract

The benefits and efficacy of control programs for herds infected with *Mycobacterium avium* subsp. *paratuberculosis* (**MAP**) have been investigated under various contexts. However, most previous research investigated paratuberculosis control programs in isolation, without modeling the potential association with other dairy diseases. This paper evaluated the benefits of MAP control programs when the herd is also affected by mastitis, a common disease causing the largest losses in dairy production. The effect of typically suggested MAP controls were estimated under the assumption that MAP infection increased the rate of clinical mastitis. We evaluated one hundred twenty three control strategies comprising various combinations of testing, culling, and hygiene, and found that the association of paratuberculosis with mastitis alters the ranking of specific MAP control programs, but only slightly alters the cost-effectiveness of particular MAP control components, as measured by the distribution of net present value of a representative U.S. dairy operation. In particular, although testing and culling for MAP resulted in a reduction in MAP incidence, that control led to lower net present value (**NPV**) per cow. When testing was used, ELISA was more cost-effective than alternative testing regimes, especially if mastitis was explicitly modeled as more likely in MAP-infected animals, but ELISA testing was only significantly associated with higher NPV if mastitis was not included in the model at all. Additional hygiene was associated with a lower NPV per cow, although it lowered MAP prevalence. Overall, the addition of an increased risk of mastitis in MAP-infected animals did not change model recommendations as much as failing to consider mastitis at all.

## 1. INTRODUCTION

Paratuberculosis, or Johne’s Disease, is a chronic intestinal disease of ruminants caused by infection with *Mycobacterium avium* subsp. *paratuberculosis* (**MAP**). Animals are usually infected at a young age, with a variable and often extended latent period [1]. Infected animals have lower milk production [2–9], decreased reproductive performance in later stages of disease [6,10–12], and are often culled early [5,13]. It is difficult to control MAP in dairy herds; many tests have poor diagnostic sensitivity [14], MAP persists in the environment for long periods of time [15], paratuberculosis symptoms are slow to develop [16], and the available vaccines are limited in distribution due to their cross-reaction with tuberculosis diagnostics [17].

The debate over the economically optimal control method for MAP results from a wide range of models and assumptions. Some studies have found test and culling to be consistently cost-effective [18,19], while others have found that cost-efficacy of test and cull required subsidized testing costs [20] or only culling of animals with decreased milk production during MAP latency [21]. Simulation models have identified cost-effective programs, such as quarterly serum enzyme-linked immunosorbent assay (**ELISA**) testing [22], quarterly milk ELISA testing [23], risk-based testing accompanied by infection control [24], vaccination or infection control [25], testing in series with ELISA and quantitative polymerase chain reaction (**qPCR**) [26], and annual fecal culture accompanied by infection control [27]. Massaro et al. [28] found that a more sensitive ELISA test could be cost-effective in US dairy herds. Others have found that hygiene improvement was effective in decreasing transmission rate [25,29], especially in combination with testing and culling [1,30]. Our previous work found that some MAP control programs were not significantly better than no control, and that some managerial practices can produce better results than some testing and culling controls (Smith et al., 2017; Verteramo Chiu et al., 2018). One factor that none of these studies addressed is the role of MAP infection in susceptibility to other infections. For example, higher mastitis incidence has been found in MAP positive farms in two different studies [32,33], and Rossi et al. [34] found that MAP-infected animals had significantly higher rates of clinical mastitis. As clinical mastitis is one of the most economically important diseases of dairy herds, a positive association between MAP infection and mastitis could greatly increase the cost-effectiveness of MAP and mastitis control. Even with no association, controlling for either disease may have spillover effects on the other disease.

The goal of this research is to examine the economic consequences of paratuberculosis in US dairy herds and the benefits of 123 specific control strategies involving various combinations of hygiene levels, types of testing, and decisions on culling, while accounting for the rise in mastitis cases associated with paratuberculosis infection.

## 2. MATERIALS AND METHODS

The infection and testing model (Figure 1) has been previously described [35], and used for an economic analysis of MAP [31]. This is a continuous-time model, simulated over 5 years after a burn-in of 50 years using values representative of US dairy herds. Details are available in the supplemental material (S1). Briefly, calves may be born susceptible or infected via vertical transmission. Susceptible calves may be infected by contact with transiently-shedding infected calves or with shedding adults. All calves age into heifers; susceptible heifers may be infected by contact with shedding adults, while infected heifers are assumed to be latently infected. All heifers age into adults. Adults infected as calves or heifers may have progressing infections, resulting in fast transition from latency, through a low-shedding phase, to high shedding and clinical disease. However, some adults infected as calves or heifers and all adults infected by contact with shedding adults experience non-progressing infections, which remain in latency for a longer period of time and only enter the low-shedding phase. All animals may be culled or die, based on an age-appropriate mortality/culling rate.

**Figure 1.**
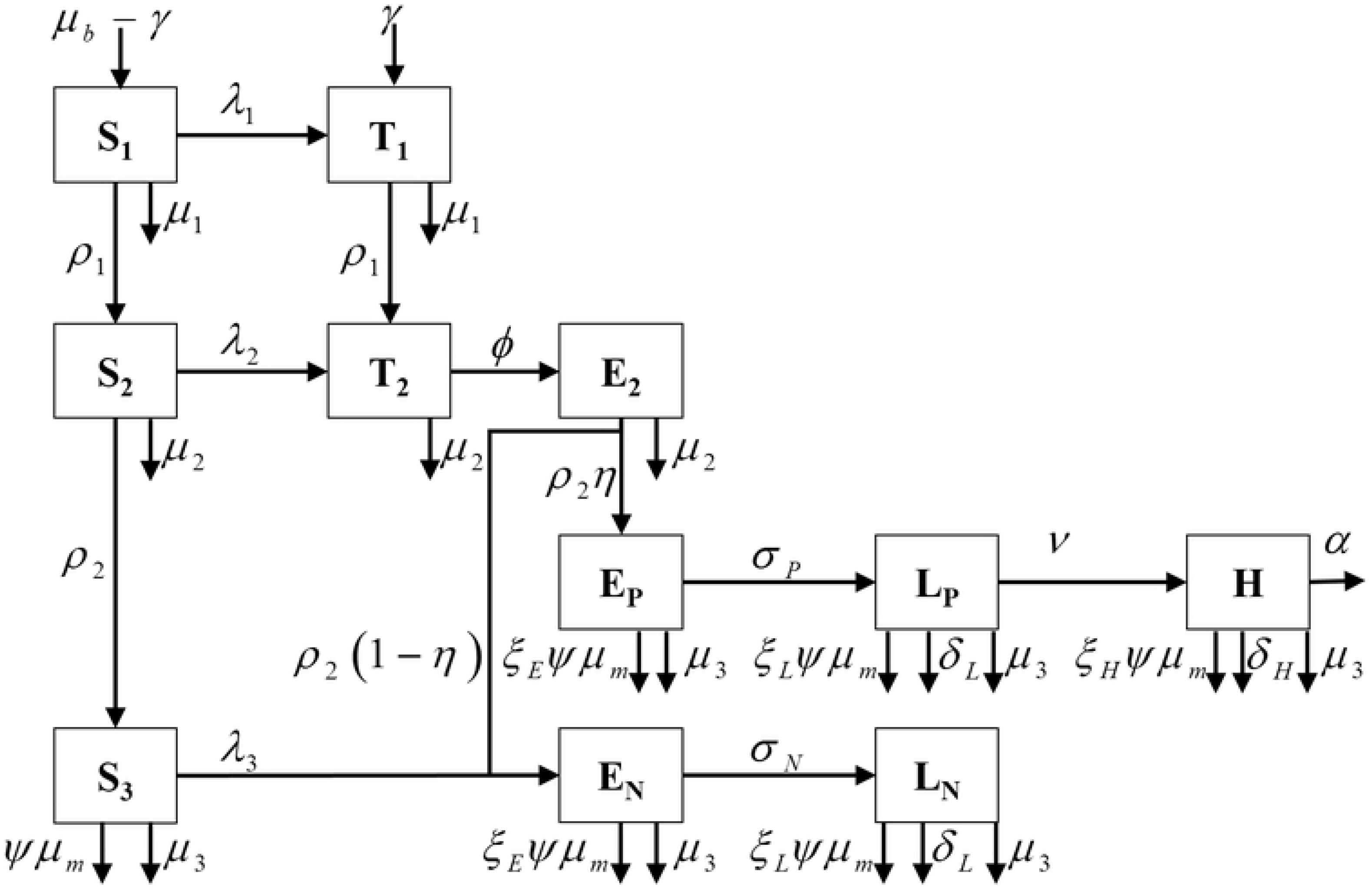
Schematic of model for *Mycobacterium avium* subsp. *paratuberculosis* in a commercial dairy herd

The economic model tracks the daily milk production of all animals in the herd and calculates the net value of the herd as the value of the milk produced plus the value of any culled animals sold, minus the cost of producing milk, the cost of raising calves, the cost of raising heifers, and the cost of MAP testing.

### 2.1 Mastitis Risk and Milk Production

The risk of first clinical mastitis (**CM**) case, and the incidence of CM, has been found to be associated with MAP infection status in cows [34], possibly due to the immune system being affected by MAP. Clinical mastitis risk was assumed to be constant for all cows; although clinical mastitis risk is known to increase with parity, this model was not age-stratified, thereby averaging out clinical mastitis risk among all animals. Annualized risk of CM by all causes was calculated from Bar et al. [36] by averaging the monthly risk over a 10 month lactation and across the first 4 lactations, then adding the total monthly risk, 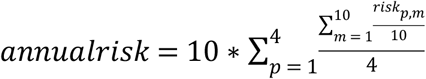. The range of possible values was identified by adding up the monthly risk for each lactation individually 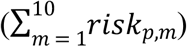. Due to the lack of data to support modeling of secondary CM cases or subclinical mastitis, these events were not modeled.

In order to determine if the effect of CM on milk production would be exacerbated by MAP status, we statistically analyzed milk production in animals with well-defined MAP infection status as described previously [9]. Briefly, we conducted a linear regression analysis to assess the effect of MAP progression (defined as progressing, non-progressing, or test-negative, where progressing animals had at least one high-positive test result) and current MAP status (defined as test-negative, latent, low-shedding, or high-shedding). In this analysis, a dichotomous term was added to indicate whether an animal had experienced a CM event in the previous 30 days. In the previous study, the linear score (log10 of the somatic cell count) was included to control for subclinical and clinical mastitis; in this analysis, that variable was not included to avoid collinearity with the CM variable.

### 2.2 Model Simulation

The model was simulated under 3 different assumptions: CM association with MAP (**MA**), no CM association with MAP (**NMA**), and no CM at all (**NM**). In the MA scenario, the rate of CM cases was assumed to be related to MAP status. The hazard ratios from the Cox proportional hazards model for MAP positive vs. negative animals [34], controlling for parity, were used to inflate the CM risk for MAP positive animals. In the NMA scenario, the rate of CM cases was assumed to be unchanged by MAP status. In the NM scenario, it was assumed that CM cases were excluded in the model. In the MA and NMA scenarios, CM occurred in susceptible adults at rate *ψ*, the annualized risk, and in MAP infected adults at rate *ξ_Iψ_*, where *ξ_I_* is the hazard ratio 1.89 [34].

Upon the occurrence of a CM case, the following actions occurred: 1) remove the milk lost due to CM, *q_mast_*, from the period’s milk production; 2) add the cost of treating CM, *t_mast_*, to the period’s cost; 3) determine if the CM case resulted in mortality. Clinical mastitis mortality was assumed to be *μ_m_*. It was assumed that no voluntary culling occurred due to a first case of CM.

The net present value (***NPV***) of each scenario was calculated as

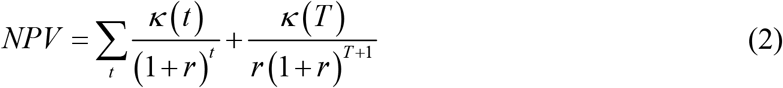

where, *t* is time in years, *κ(t)* is the value of the herd in year *t, r* is the discount rate, and T is the final time period. The second term represents a terminal wealth term, the NPV of the last year cash flow continuing into perpetuity, to account for the value of the herd going forward past the terminal year of T. Herds were simulated for 5 years, which is considered a realistic planning window for commercial dairy herds.

### 2.3 Determining Stochastic Dominance

Ranking of control programs by the distribution of NPV from 100 iterations of a five year period was performed using first and second-order dominance [37]. First-order and second-order stochastic dominance are methods of determining preference for an activity with variable (stochastic) results; a dominant strategy by either method is to be preferred to its comparator. First-order stochastic dominance is relevant for decision makers who prefer more wealth to less wealth (increasing utility function), and second-order stochastic dominance is relevant for decision makers who in addition to preferring more to less wealth are also risk averse (increasing and concave utility function).

Briefly, if NPV_A_ is the cumulative distribution function of the NPV of control strategy A and NPV_B_ is the cumulative distribution function of the NPV of control strategy B, first-order dominance of strategy A states that

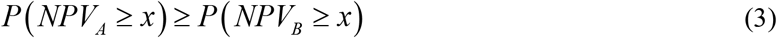

for all possible values of *x* (the range of simulated NPV values for a farm in a given scenario) and with a strict inequality for at least one value of x.

Likewise, second-order dominance of strategy A states that

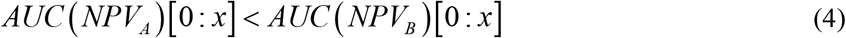

for all possible values of *x* (as in equation 3), where *AUC(i)[0:x]* is the area under the curve of the cumulative distribution function of distribution *i* from 0 to *x*.

### 2.4 Analyzing Dominance Results

Each of the 123 control strategies comprises four components: hygiene level, test used, test frequency, and which animals are culled (Supplemental Information S2). To estimate the effect of each of these components on dominance, we estimated a linear regression of the proportion of dominated strategies under SOSD on each of the strategies’ components, measured by dummy variables. The econometric model has the following form,

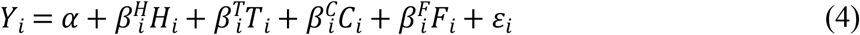

Where *Y* is the proportion that strategy *i* SOSD the other strategies, where the value of *Y* ranges from 0 to 1. *H* is an n × 2 matrix of hygiene level indicators (standard and high hygiene), *T* is an n × 4 matrix of test indicator (FC, ELISA, PCR, and hypothetical testing of calves with FC), *C* is an n × 2 matrix of culling policy (cull all test positive, and cull all high shedders), *F* is an n × 4 matrix of frequency of testing (annual, biannual, continuous annual, continuous biannual). Parameter *α* is the intercept term, which includes the effects of moderate hygiene, and culling after 2 positive tests. Parameters *β^H^, β^T^, β^C^, β^F^* are vectors to be estimated for each of the components of the strategies. This regression is conducted separately for each of the three mastitis assumptions and assumes additive linear effects only. To determine which factors are most associated with changes in the NPV, this analysis was repeated using the difference between the NPV of a particular iteration and the NPV of no control from the same starting condition, expressed as an amount per cow, as the *Y*. Each of these analyses was repeated for each of the mastitis assumptions (NM, NMA, and MA) and the fitted coefficients were compared. The analysis was repeated at two different herd sizes (100 and 1000) and two different initial MAP prevalence levels (7% and 20%), but results will focus on the 1000-head herd with 20% initial prevalence.

### 2.5 Sensitivity Analysis

Global sensitivity analysis was performed using optimized Latin Hypercube sampling via the *lhs* package [39] with 500 parameter sets. For each parameter set, a 1000 head herd was simulated 100 times from the same randomly drawn initial population values under three generalized culling strategies (none, cull all positive adults, and cull all positive calves) with and without improved hygiene. Impact of parameters on NPV was determined using the Pearson’s rank correlation coefficient. All parameters used are shown in Tables 1–3. Where variability in parameters was not provided by the source, parameters were varied by ± 10% for the sensitivity analysis. Where variability in parameters was available, parameters were varied over their interquartile ranges. Testing parameters were varied over the range of the interquartile ranges of all tests, and hygiene parameters were varied over the range of the possible additional hygiene levels. Parameters were considered significantly related to NPV at the level of α = 0.05 with Bonferroni’s correction.

**Table 1.** Biological parameters and interquartile ranges (IQR) used in a model of *Mycobacterium avium* subsp. *paratuberculosis* and clinical mastitis co-infection in a dairy herd.

| Par. | Description | Value (IQR) | Source |
| --- | --- | --- | --- |
| $\mu_1$ | removal rate of calves (/year) | 0.09 (0.08-0.1) | [40] |
| $\mu_2$ | removal rate of heifers (/year) | 0.01 (0.008-0.015) | [40] |
| $\mu_3$ | removal rate of adults (/year) | 0.35 (0.3-0.4) | [40] |
| $\mu_{b,base}$ | birth rate of female calves (/adult/year) | 0.45 (0.4-0.5) | [40] |
| $\mu_{b,H}$ | birth rate of high shedding dams (/adult/year) | 0.15 (0.1-0.45) | [11] |
| $\gamma_E$ | proportion of calves of latent animals infected at birth | 0.01 (0-0.04) | [40] |
| $\gamma_L, \gamma_H$ | proportion of calves of shedding animals infected at birth | 0.04 (0.01-0.08) | [40] |
| $\rho_1, \rho_2$ | aging rate (/year) | 1 (0.8-1.2) | Assumed |
| $\eta$ | proportion of infected heifers becoming progressing adults | 0.335 (0.5-1) | [40] |
| $\varphi$ | transition rate from transient shedding to latent (/year) | 2 (0.8-3) | [40] |
| $\sigma_L$ | transition rate from latent to low shedding, low path (/year) | 0.53 (0.44-0.67) | [35] |
| $\sigma_H$ | transition rate from latent to low shedding, high path (/year) | 21.5 (1.75-40) | [35] |
| $\nu_H$ | transition rate from low to high shedding, high path (/year) | 1.08 (0.75-1.94) | [35] |
| $\alpha$ | clinical disease-related culling rate (/year) | 0.67 (0.5-0.8) | [11] |
| $\beta$ | transmission coefficient for 7% and 20% initial prevalence (/year) in a 1000-head herd | 0.001, 0.003 (0.0023-0.0012) | calculated |
| $e_\beta$ | proportional transmission effect due to improved and moderately improved hygiene | 0.6, 0.98 (0.6-0.98) | [41,42]<br>[43] |
| $\psi$ | risk of clinical mastitis (/year) | 0.27 (0.12-0.42) | [36] |
| $\xi_I$ | hazard ratio for clinical mastitis in infected cows | 1.89 (1.53-2.33) | [34] |
| $\mu_m$ | risk of mortality during clinical mastitis | 0.0175 (0.01-0.02) | [36] |

**Table 2.**
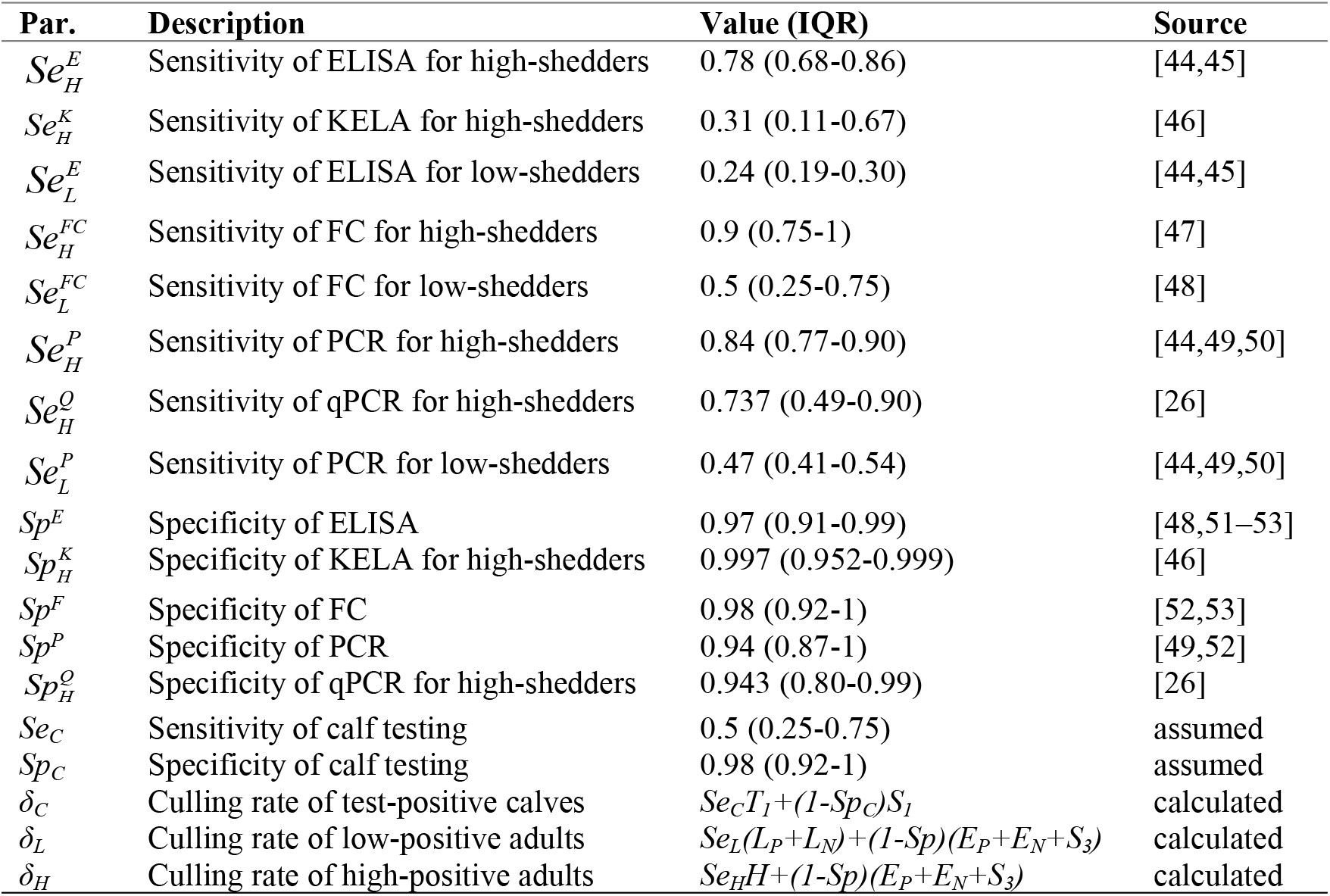
Testing parameters and interquartile ranges (IQR) used in a model of *Mycobacterium avium* subsp. *paratuberculosis* infection in a dairy herd.

| Par. | Description | Value (IQR) | Source |
| --- | --- | --- | --- |
| $Se_H^E$ | Sensitivity of ELISA for high-shedders | 0.78 (0.68-0.86) | [44,45] |
| $Se_H^K$ | Sensitivity of KELA for high-shedders | 0.31 (0.11-0.67) | [46] |
| $Se_L^E$ | Sensitivity of ELISA for low-shedders | 0.24 (0.19-0.30) | [44,45] |
| $Se_H^{FC}$ | Sensitivity of FC for high-shedders | 0.9 (0.75-1) | [47] |
| $Se_L^{FC}$ | Sensitivity of FC for low-shedders | 0.5 (0.25-0.75) | [48] |
| $Se_H^P$ | Sensitivity of PCR for high-shedders | 0.84 (0.77-0.90) | [44,49,50] |
| $Se_H^Q$ | Sensitivity of qPCR for high-shedders | 0.737 (0.49-0.90) | [26] |
| $Se_L^P$ | Sensitivity of PCR for low-shedders | 0.47 (0.41-0.54) | [44,49,50] |
| $Sp^E$ | Specificity of ELISA | 0.97 (0.91-0.99) | [48,51-53] |
| $Sp_H^K$ | Specificity of KELA for high-shedders | 0.997 (0.952-0.999) | [46] |
| $Sp^F$ | Specificity of FC | 0.98 (0.92-1) | [52,53] |
| $Sp^P$ | Specificity of PCR | 0.94 (0.87-1) | [49,52] |
| $Sp_H^Q$ | Specificity of qPCR for high-shedders | 0.943 (0.80-0.99) | [26] |
| $Se_C$ | Sensitivity of calf testing | 0.5 (0.25-0.75) | assumed |
| $Sp_C$ | Specificity of calf testing | 0.98 (0.92-1) | assumed |
| $\delta_C$ | Culling rate of test-positive calves | $Se_C T_1 + (1 - Sp_C) S_1$ | calculated |
| $\delta_L$ | Culling rate of low-positive adults | $Se_L (L_P + L_N) + (1 - Sp) (E_P + E_N + S_3)$ | calculated |
| $\delta_H$ | Culling rate of high-positive adults | $Se_H H + (1 - Sp) (E_P + E_N + S_3)$ | calculated |

**Table 3.** Economic parameters and inter-quartile ranges (IQR) used in a model of *Mycobacterium avium* subsp. *paratuberculosis* and mastitis co-infection in dairy herds.

| Cost | Description | Value (IQR) | Reference |
| --- | --- | --- | --- |
| $prev$ | prevalence of MAP infection in purchased cows | 0.094 (0.077-0.111) | [54] |
| $C_{FC}$ | cost of fecal culture test per animal | \$36 (\$25-\$42) | [55–58] |
| $C_E$ | cost of ELISA test per animal | \$6 (\$4-\$8) | [55–59] |
| $C_P$ | cost of PCR test per animal | \$32 (\$32-40) | [55–59] |
| $\omega_{hyg,mod}$ | Annual cost of implementing moderate hygiene per adult (clean milk) | \$35.54 | [21] |
| $\omega_{hyg,large}$ | Annual cost of implementing improved hygiene per adult (clean milk, separate calving pens, separate housing) | \$49.64 | [21] |
| $C_{cow}$ | Daily operating cost per kg milk produced | \$0.35 (0.33-0.37) | [60] |
| $C_{heifer}$ | Daily operating cost of raising a calf/heifer | \$2.995 (2.662-3.403) | [61] |
| $P_{cow}$ | Cull-cow price per kg | \$1.9671 (1.7292-2.31) | [62] |
| $P_{milk}$ | Milk price per kg | \$0.444 (0.394-0.482) | [62] |
| $P_{sale}$ | Sale price of replacement heifer | \$2232 (2000-2500) | [61] |
| $Q_{cull}$ | Average cull cow weight | 680.4 kg (660-700) | [63] |
| $Q_{milk,S}$ | Average daily milk production per uninfected cow | 32.62 kg (25.45-42.73) | [9] |
| $Q_{milk,EN}$ | Average daily milk production per latent cow (non-progressing) | 32.17 kg (24.09-42.73) | [9] |
| $Q_{milk,LN}$ | Average daily milk production per low-shedding cow (non-progressing) | 30.94 kg (24.09-42.73) | [9] |
| $Q_{milk,EP}$ | Average daily milk production per latent cow (progressing) | 33.12 kg (22.27-39.09) | [9] |
| $Q_{milk,LP}$ | Average daily milk production per low-shedding cow (progressing) | 29.13 kg (22.27-39.09) | [9] |
| $Q_{milk,H}$ | Average daily milk production per high-shedding cow | 22.17 kg (12.27-32.27) | [9] |
| $\psi_H$ | Proportional adjustment in cull weight for high-shedding cows | 0.9 (0.75-1) | assumed |
| $t_{mast}$ | Treatment cost per clinical mastitis case | \$50 (35.50-73.50) | [36] |
| $q_{mast}$ | Milk loss per clinical mastitis case | 90.3 kg (64-183) | [64] |
| $r$ | Discount rate | 0.02 (0.01-0.08) | assumed |

## 3. RESULTS

### 3.1 Milk Production Results

There were 31,583 monthly milk observations available for analysis, of which 537 occurred within a month of a CM event. Of those, 424 were in test-negative individuals, 97 were in non-progressing animals (85 latent, 12 low-shedding), and 16 were in progressing animals (14 latent, 2 low-shedding). Adding a variable indicating a recent CM did not improve the fit of the model including an interaction between MAP progression and status (BIC=216,536 with the term and BIC=216,527 without the term), so milk loss in animals with both MAP infection and CM was simulated to be additive.

### 3.2 Model Results

For simplicity, we will present rankings of only 13 potential control options, and the results of SOSD only, as SOSD implies FOSD. The base for comparison is no MAP control. If testing is used, we assume that it will be based on the serum enzyme-linked immunosorbent assay (ELISA) test, administered to all animals either annually or biannually. Animals may be culled after any positive test, only after a test result indicating a high-positive response, or only after the second positive test result. Additionally, the farm may choose to continue ELISA testing after 5 negative whole-herd tests, or to discontinue testing after the 5th negative whole-herd test.

Model results for this subset of control options are shown in Table 4 for a 1,000 head herd with 20% initial shedding prevalence under CM association with MAP (**MA**), no CM association with MAP (**NMA**), and no CM at all (**NM**); results for other initial herd sizes and initial shedding prevalence are generally similar, and are shown in Supplemental Table S3. The model predicted that all control programs would decrease the median true infection prevalence of paratuberculosis over 5 years, and most would decrease the median shedding prevalence (Figure 2).

**Table 4.**
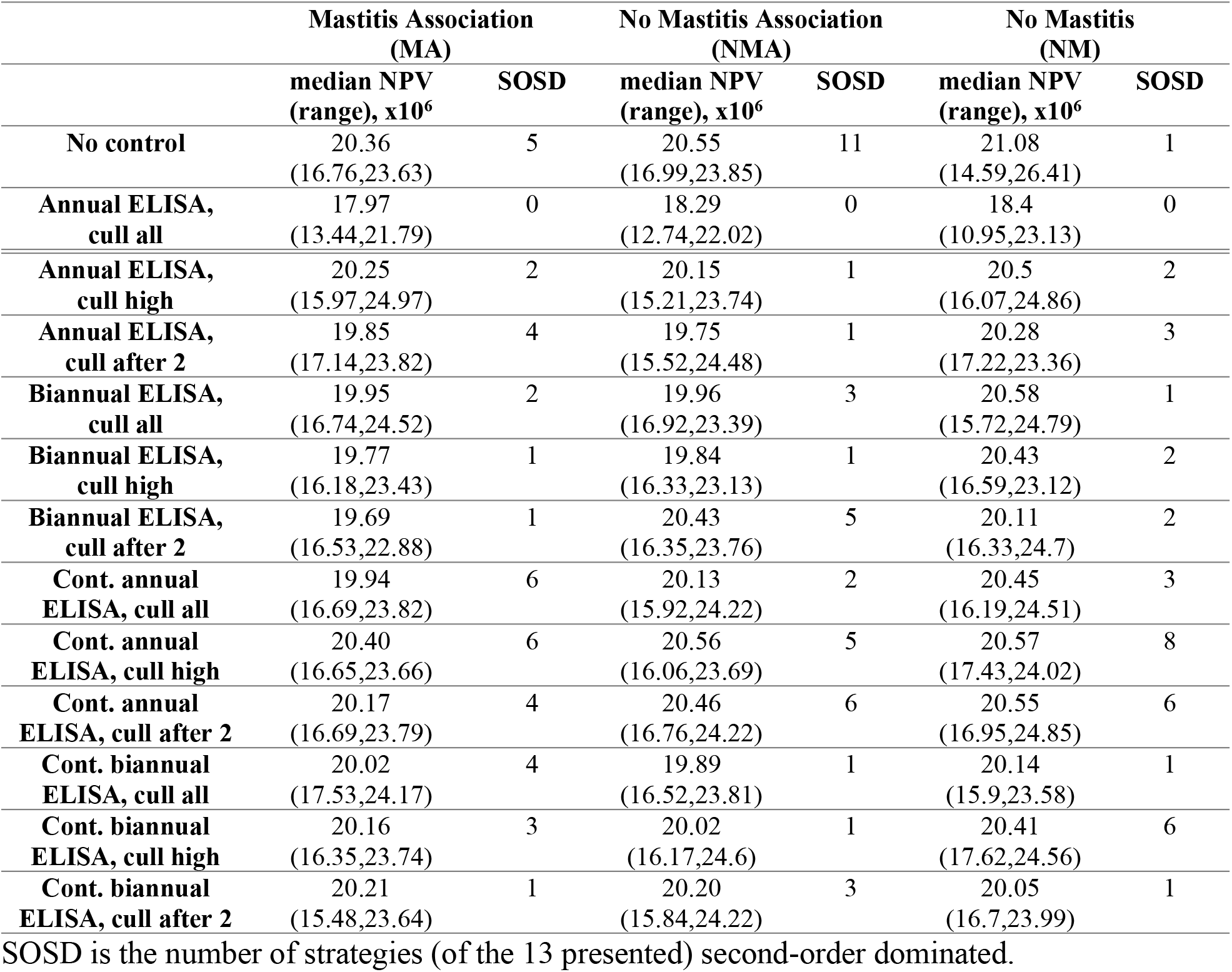
ELISA-based testing strategies and their NPV distribution and number of second-order dominated strategies for each mastitis scenario. Results are for a 20% initial MAP prevalence in a 1,000 head herd.

**Figure 2.**
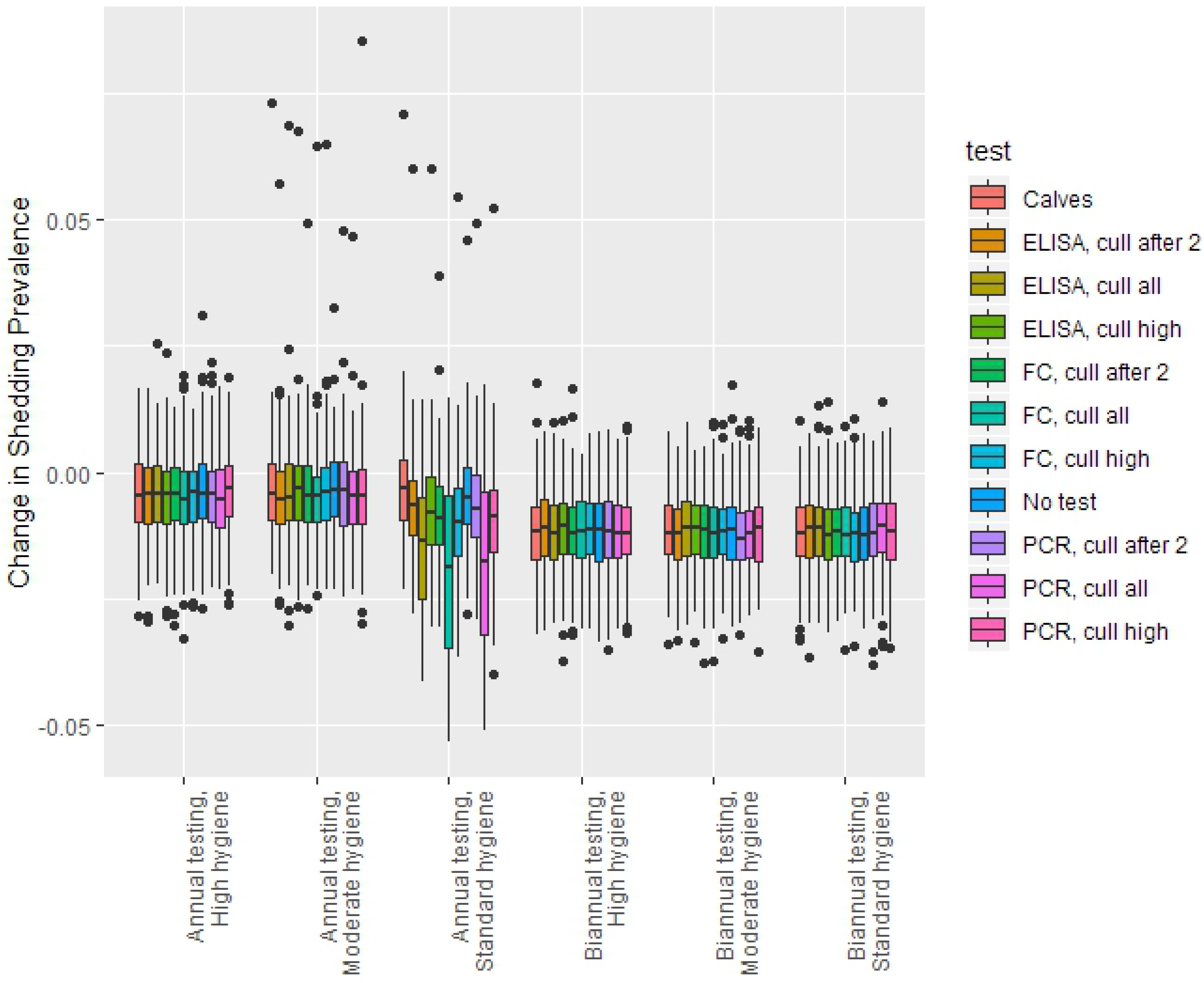
Predicted change in shedding prevalence of paratuberculosis infection after 5 years of control in a 1,000 cow herd with a median initial prevalence of 20%.

### 3.3 Stochastic Dominance Results

Regression results for the econometric model of SOSD rank for each of the CM scenarios and MAP prevalence are shown in Figure 3 and Table 5, using results of all 123 control combinations (S2). All six regressions have a similar set of significant variables, but their effects can be different across herds. Not testing was consistently significantly worse than annual testing, with the exception being in a herd with low initial prevalence and assuming NMA. Biannual testing was not significantly different from annual testing in most scenarios. Continuing to test after 5 negative whole-herd tests was significantly better than discontinuing testing. ELISA testing was significantly better than FC or testing calves in the MA and NM scenarios, but not in the NMA scenario. However, in the NMA scenario with high initial prevalence, FC and PCR were significantly worse than testing calves. High levels of hygiene were significantly worse than standard in all cases, and moderate levels of hygiene were significantly worse than standard in most cases.

**Figure 3.**
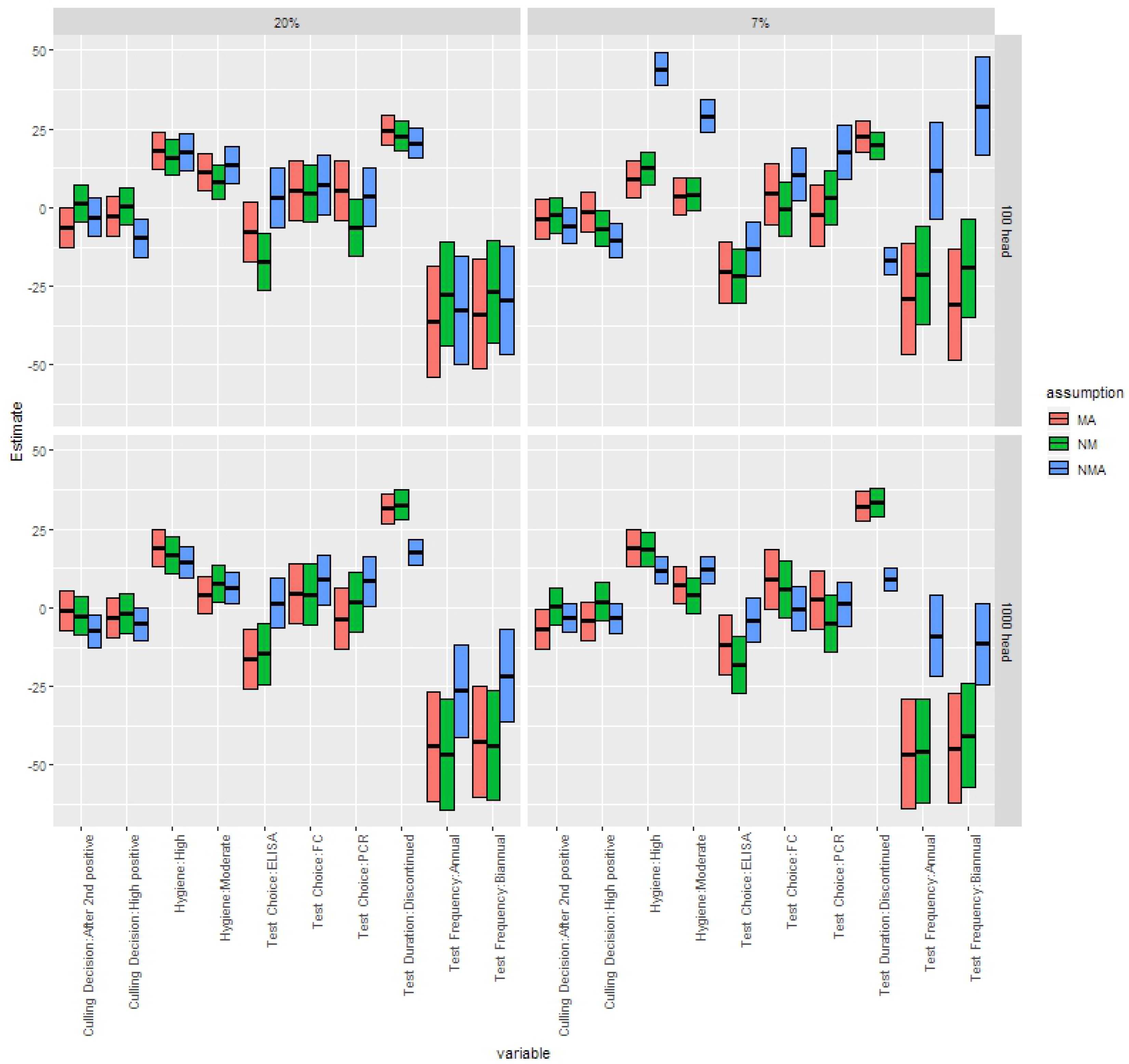
Coefficients from multivariable linear regressions for the overall second-order stochastic dominance rank (1=best, 123=worst) of MAP control programs, separated by herd size, initial shedding prevalence, and assumption about relationship between MAP and clinical mastitis (MA: mastitis association; NMA: no mastitis association; NM: no mastitis). Central bar is estimate, box shows 95% confidence interval around estimate.

**Table 5.**
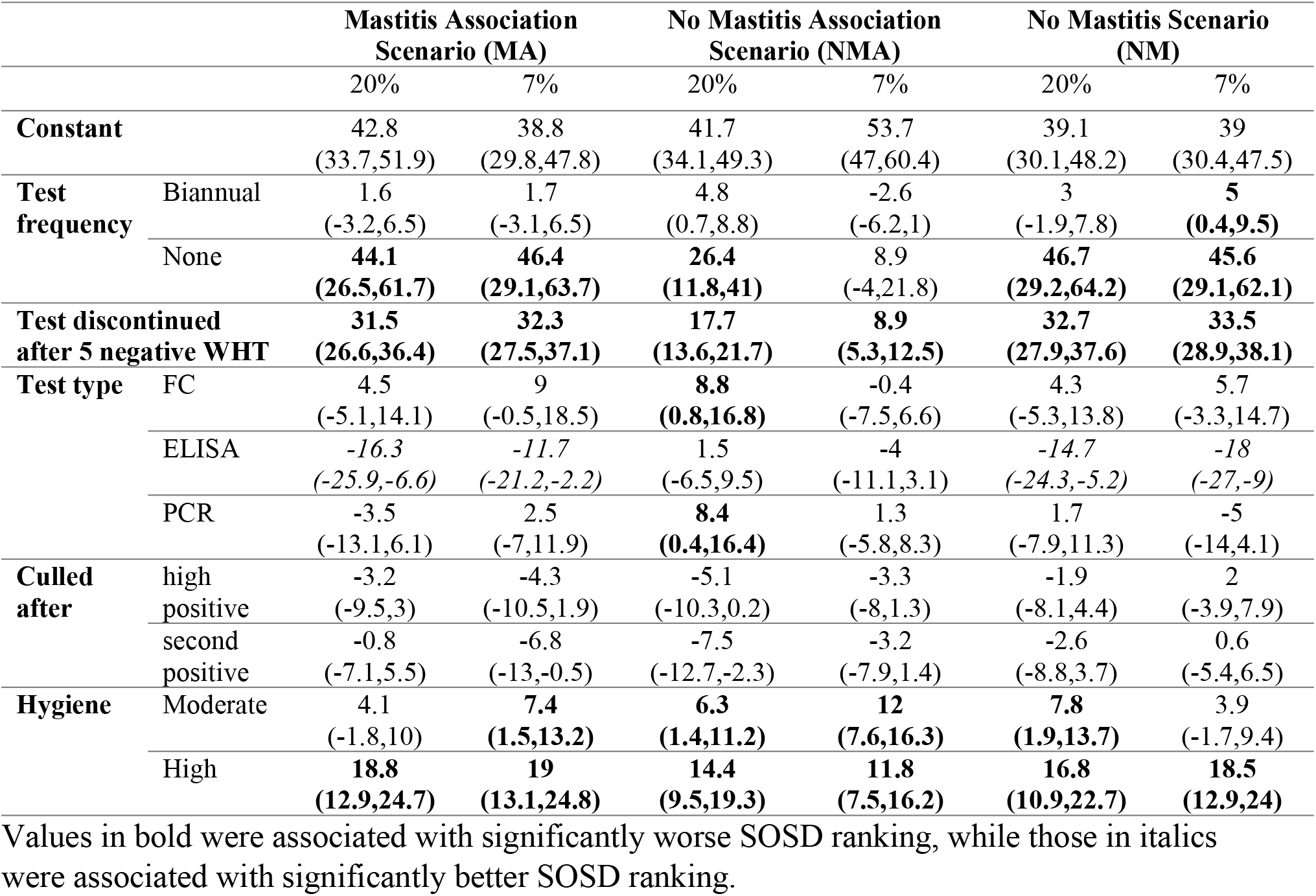
Linear regression results of the proportion of dominated strategies under SOSD on strategy characteristics on all initial herds. Constant term includes the effects of annual continuous testing of calves, culling animals after one positive test, and standard hygiene. Model assumes additive linear effects.

Regression results for the econometric model of NPV for each of the CM scenarios and herds are shown in Figure 4. Biannual testing was significantly associated with a lower NPV compared to not testing, as was annual testing in almost all cases. Use of an ELISA test was significantly associated with a higher NPV than other test choices. Moderate or high hygiene levels were significantly associated with a lower NPV than standard hygiene. In a large herd, culling only high positive cows or after the second positive test was significantly associated with a slightly higher NPV; this relationship was not seen in small herds. There were few differences in coefficient values across CM scenarios.

**Figure 4.**
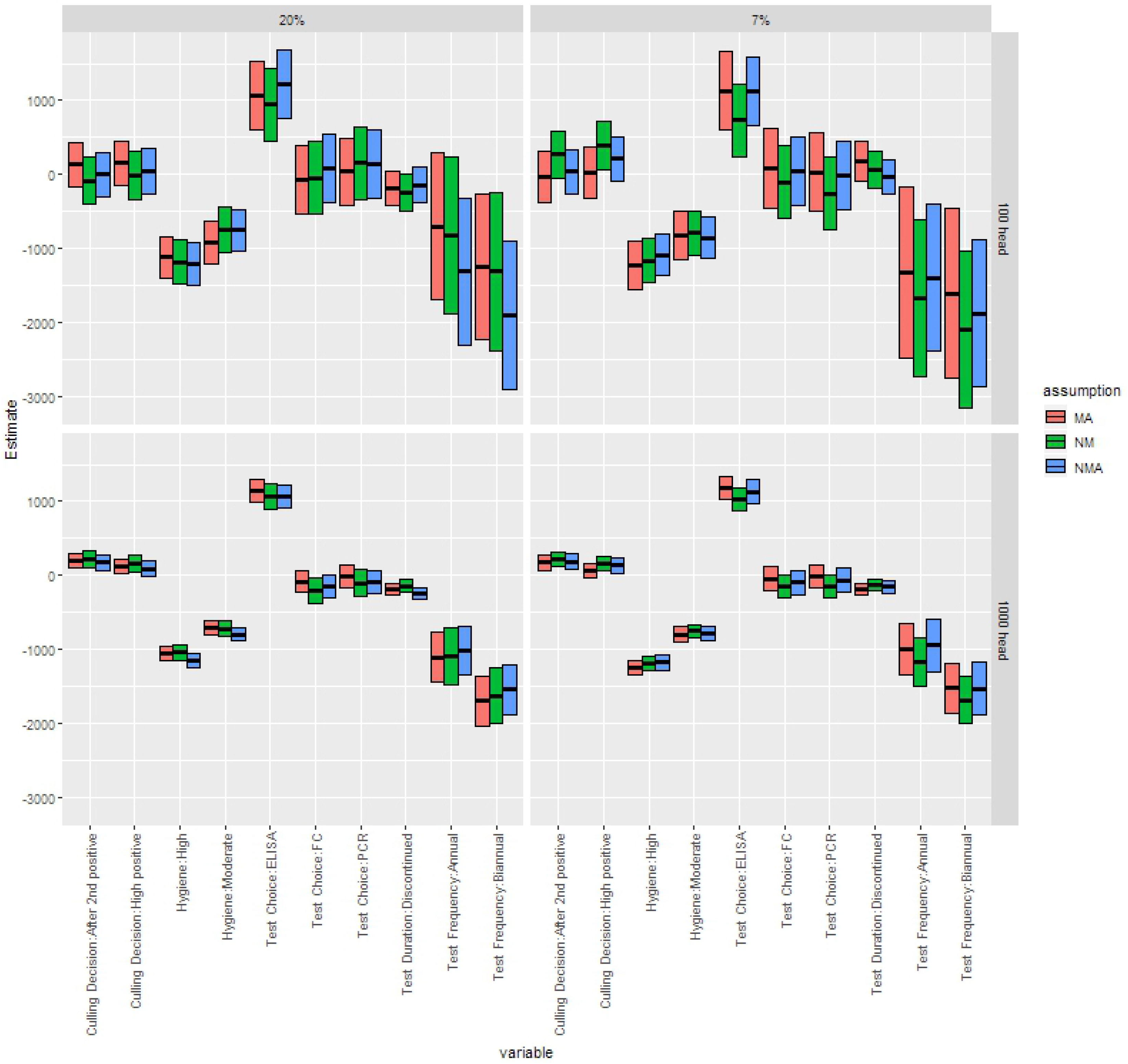
Coefficients from multivariable linear regressions for change in NPV per cow by adding MAP control programs, separated by herd size, initial shedding prevalence, and assumption about relationship between MAP and clinical mastitis (MA: mastitis association; NMA: no mastitis association; NM: no mastitis). Central bar is estimate, box shows 95% confidence interval around estimate.

### 3.4 Sensitivity Analysis Results

The partial rank correlation coefficients of all significantly correlated parameters from the global sensitivity analysis are shown in Figure 5. Most scenarios had the same parameters consistently related with NPV, primarily economic and production-related parameters. The risk of clinical mastitis was significantly related to NPV in all scenarios.

**Figure 5.**
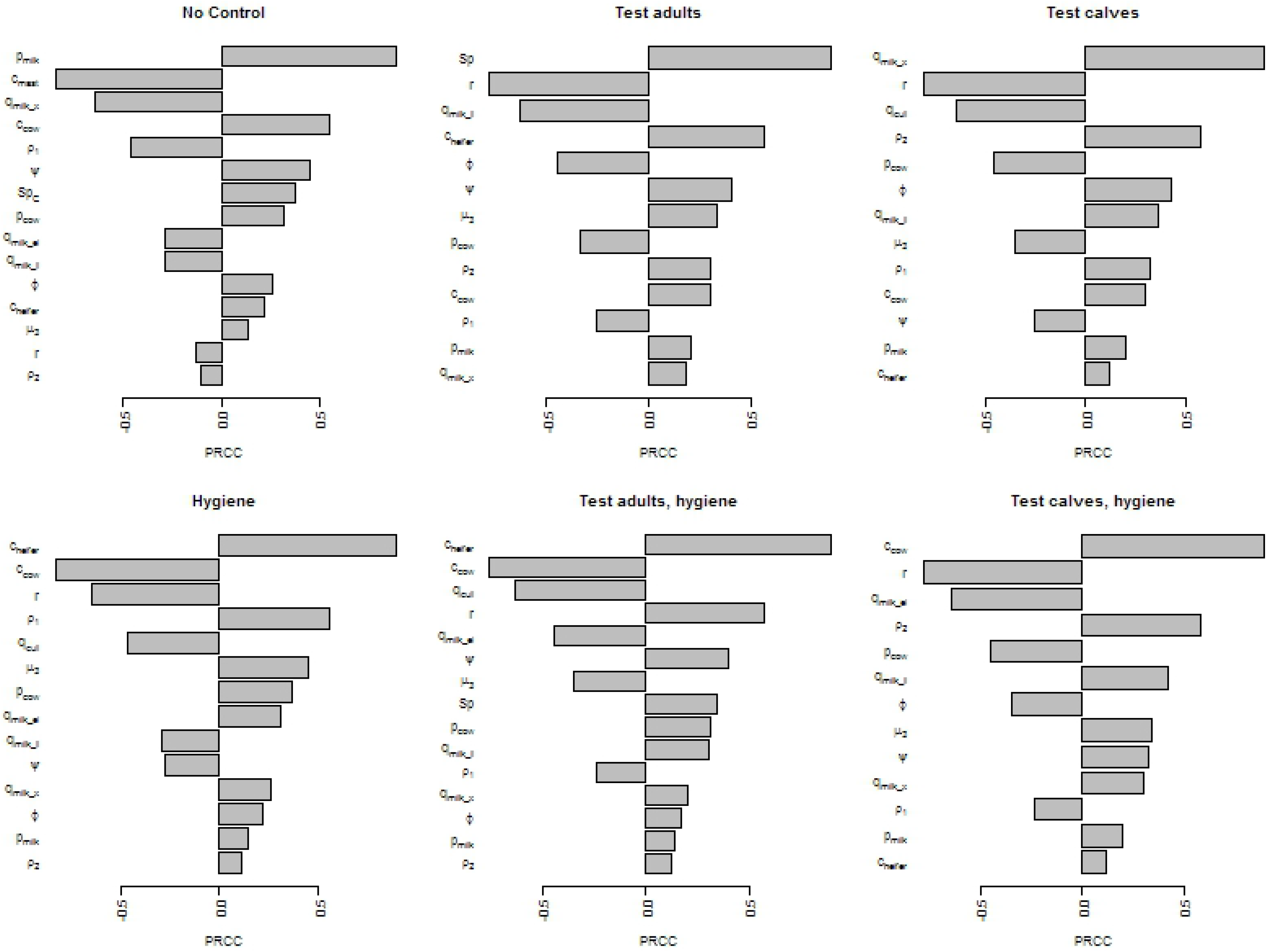
Partial rank correlation coefficients (PRCC) from a global sensitivity analysis on net present value over 5 years of paratuberculosis control, assuming an association between mastitis incidence and MAP infection.

## 4. DISCUSSION

This research shows that consideration of interacting disease systems does not importantly change the results of an economic analysis of disease control. Adding an increased rate of CM among infected animals to an economic model of paratuberculosis control only slightly changed the ranking of control programs. Specifically, failing to include CM in the model resulted in a weaker preference for standard hygiene alone. Including CM but not its association with paratuberculosis resulted in a stronger preference for standard hygiene alone, biannual ELISA testing and culling adults after 2 positive tests, and continuous ELISA with the same culling policy. Culling for paratuberculosis should in theory have the side benefit of partially controlling for CM. However, the inclusion of an association between CM and paratuberculosis did not change the overall conclusions of this economic model. This is likely due to two factors: the high cost of MAP control and the relatively small size of the impact of MAP on mastitis.

We believe that the high cost of MAP control is the reason that few control strategies have been shown to economically dominate no control. If the cost of implementing testing or hygiene, not including costs related to culling of animals, were removed from the NPV, the distributions are somewhat similar for many control programs (Supplemental Figure, S5). However, the cost of these programs is high (Supplemental Table, S4): over a 5 year period, in a 1,000-head herd, the discounted cost of testing all adults annually via ELISA was calculated at $16,148. Testing all adult cows biannually using fecal culture or PCR was more than an order of magnitude higher. These numbers do not include the costs of culling test-positive animals, or the lower income due to smaller milking herd sizes after test-based culling in closed herds, each of which would raise the cost of control even more.

Previous studies have disagreed as to the cost-effectiveness of testing for MAP. While some models suggest that test and cull programs are effective at reducing the prevalence of MAP [30], others suggest that they are not sufficient to control MAP by themselves [23,65]. Our work here has shown that they are capable of decreasing the shedding prevalence of MAP, but are unlikely to be cost-effective. The exception would be ELISA testing, which others have also found to be potentially cost-effective [28]. This is likely due to the low cost and fast turn-around time for ELISA results.

We found here that hygiene was not cost-effective by any measure, and that this was unrelated to the relationship between MAP and mastitis. We had hypothesized that expensive control programs such as hygiene improvement (estimated here to cost a 1,000-head herd between $95,652 and $133,600 over a five year period) would become cost-effective as their effect on other pathogens was considered. The hygiene changes made to improve MAP control, however, are unlikely to directly impact CM incidence. While other models have suggested that hygiene changes are indeed cost-effective [29], these may be assuming a lower base hygiene level than our simulated herds. There also may be more benefits from hygiene improvement over a longer time frame than the 5 years used here.

Our model did not show a strong effect of paratuberculosis association with CM on economically optimal control. The global sensitivity analysis also showed that the hazard ratio for CM incidence in MAP-infected animals was not significantly associated with NPV. Likely, this is because the association between paratuberculosis and CM is so small in a practical sense. The hazard ratio for first CM cases among MAP-infected animals is 1.89 (IQR: 1.53-2.33). However, with an annualized rate of 0.27 CM cases/animal/year, this translates into an annual average of 27 extra cases of CM in a 1,000 head herd with 20% MAP infection prevalence. Given a cost per case of CM of $90, not counting mortality, the additional cost to the herd is approximately $2,500. Discounted over a 5 year simulation period, that results in a total cost of $11,655 due to additional CM cases. This is less than the cost of the least expensive MAP control program (annual ELISA testing), and, as no program can immediately eliminate MAP in the herd, not all of the potential cost from the increased CM cases would be avoided by implementing control.

One large limitation of this model was the lack of age stratification, resulting in the necessary simplification of constant clinical mastitis risk. It is known that clinical mastitis risk increases with parity [66] and changes throughout the lactation [36]. However, accounting for age and lactation stage in a compartmental model would cause the model to become intractable. For more realistic modeling frameworks, it becomes necessary to transition to a more computationally demanding modeling system, such as the agent-based model presented in Verteramo Chiu et al. [67].

Regardless of the effects of MAP associations with CM, some overall preferences were determined. On average, continuing to test and cull after 5 negative whole-herd tests was always preferred. ELISA was the best-ranked test, followed by no testing. Standard hygiene was always preferred, with increasing hygiene levels associated with economically worse-ranked programs.

## 5. CONCLUSION

We have found that, in the setting of a typical commercial US dairy, the addition of clinical mastitis to a model for MAP control only slightly changed the ranking of individual control programs, but did not greatly change the overall cost-effectiveness of components of MAP control. These suggest that only testing by ELISA may be cost-effective.

## 6. ACKNOWLEDGEMENTS

The authors gratefully acknowledge funding provided by the National Institute of Food and Agriculture of the United States Department of Agriculture through NIFA Award No. 2014-67015-2240 as part of the joint USDA-NSF-NIH-BBSRC-BSF Ecology and Evolution of Infectious Diseases program. The funding sources played no role in the research.

## Supporting Information Captions

**S1 Table: Events, changes, and rates used for simulation via Gillespie’s direct algorithm**

**S2 Table: Paratuberculosis control strategy IDs**

**S3 Table: ELISA-based strategies and their NPV distribution and number of dominated strategies for each mastitis scenario and herd type.**

**S4 Table: Discounted cost of implementing different possible paratuberculosis controls, not including culling and replacement costs, over a 5 year period in a 1,000-head dairy herd**

**S5 Figure: Net Present Value and discounted cost of control for each paratuberculosis control strategy over 5 years in a 1,000-head dairy herd with 7% initial paratuberculosis prevalence and increased mastitis in paratuberculosis-infected cows**

